# SpliceV: Analysis and publication quality printing of linear and circular RNA splicing, expression and regulation

**DOI:** 10.1101/509661

**Authors:** Nathan Ungerleider, Erik Flemington

## Abstract

In eukaryotes, most genes code for multiple transcript isoforms that are generated through the complex and tightly regulated process of RNA splicing. Despite arising from identical precursor transcripts, alternatively spliced RNAs can have dramatically different functions. Transcriptome complexity is elevated further by the production of circular RNAs (circRNAs), another class of mature RNA that results from the splicing of a downstream splice donor to an upstream splice acceptor. While there has been a rapid expansion of circRNA catalogs in the last few years through the utilization of next generation sequencing approaches, our understanding of the mechanisms and regulation of circular RNA biogenesis, the impact that circRNA generation has on parental transcript processing, and the functions carried out by circular RNAs remains limited. Here, we present a visualization and analysis tool, SpliceV, that rapidly plots all relevant forward-and back-splice data, with exon and single nucleotide level coverage information from RNA-seq experiments in a publication quality format. SpliceV also integrates analysis features that assist investigations into splicing regulation and transcript functions through the display of predicted RNA binding protein sites and the configuration of repetitive elements along the primary transcript. SpliceV is compatible with both Python 2.7 and 3+, and is distributed under the GNU Public License. The source code is freely available for download at https://github.com/flemingtonlab/SpliceV and can be installed from PyPI using pip.

## Background

The majority of mammalian genes code for multiple transcript isoforms that contribute substantially to the vast complexity of both the mammalian transcriptome and proteome^1,2^. Each mature isoform is generated through a dynamic series of tightly coordinated actions that begin to occur as the nascent transcript is being synthesized^3^. The growing precursor RNA is sequentially bound by a myriad of RNA binding proteins (RNABPs) and small nucleolar RNAs (snoRNAs; reviewed in *Wahl et al*^*4*^) as the exon-intron boundaries become defined through these specific ribonucleoprotein complex interactions. The assembled ribonucleoprotein complex, termed the spliceosome, facilitates intron excision and covalent ligation of flanking exons across the gene locus, ultimately generating a mature transcript isoform.

While each exon-intron boundary inherently contains a splice site, contiguous exons are not always spliced together. Retained introns^5^, skipped exons^6^, and cryptic splice sites^7^ commonly diversify the profile of fully processed transcript isoforms. Splice site proximity, defined by RNA secondary structure, is a major factor in splice site selection^8^. Intron length and the presence or absence of inverted repeats can impact the physical distance between splice donor and acceptor^9^. Branch point sequence motifs^10^ and nucleotides adjacent to splice sites^11^ fine tune the strength of snoRNA interactions. Further, variations in polypyrimidine tracts can preferentially attract one RNABP over another^12^. An additional layer of regulation is provided by the cellular abundance and availability of individual RNABPs and snoRNAs, allowing for tissue and context specificity of RNA processing and alternative isoform expression^13^.

Circular RNAs (circRNAs) are covalently closed circular transcripts that are generated through the splicing of downstream splice donors to upstream splice acceptors by the spliceosome, frequently utilizing canonical forward-splice junction sequences^14^. This class of RNA has recently been shown to be evolutionarily conserved^15,16^, highly abundant in humans^15^, and for many genes represents the most prevalent isoform^15^. The 3’ to 5’ backsplicing reaction, required for circRNA biogenesis, correlates with the speed of precursor transcript elongation^17^, occurs more frequently at splice sites flanked by long introns^18^ and introns containing reverse complementary sequences^19^. To date, little is known regarding the function of the vast majority of circRNAs. Nevertheless, some circRNAs have been shown to function as microRNA sponges^20,21^, as direct regulators of parental gene expression^22^, in signaling between cells^23^ and as templates for translation^24,25^.

The primary output of gene expression analysis pipelines is a dataframe with one value assigned to each gene. Individual gene expression values are calculated by tallying up reads that align to any region along each gene, followed by normalizing to the length of the gene and correcting for the total number of aligned reads in that sample. While useful for broad scale interpretations, these gene level expression values fail to resolve the abundances of individual isoforms of each gene, the function of which can dramatically differ from one another. Importantly, alternatively spliced isoforms can code for proteins that are truncated^26^, lack specific functional domains^27^, are completely unique^28^, and in some cases, alter cell fate entirely^26,29^. Here we present a visualization tool, SpliceV, that facilitates detailed exploration and visualization of transcript isoform expression in publication quality format. SpliceV facilitates within-and across-sample analyses and includes the display of predicted cis and trans regulatory factors to further assist in the biogenesis and function studies. Together, SpliceV should be a useful tool for a wide spectrum of the RNA biology research community.

## Implementation

Our software package is written in Python 3 but is backwards compatible with Python 2.7, relying only upon the third-party libraries, matplotlib^30^, numpy^31^, and pysam^32^. Source code can be found at https://github.com/flemingtonlab/SpliceV and can be installed from PyPI using the Python package manager, pip. SpliceV is written with a GNU 3.0 public license, provided with anonymous download and installation. Full usage information can be found in Supplementary File 1.

SpliceV generates plots of coverage, splice junctions, and back-splice junctions with customizable parameters, depicting expression of both the linear and circular isoforms of a given gene. Standard formats (BAM and GTF) are accepted as input files, and an unlimited number of samples can be analyzed and compared at once. Because junction calling sensitivity can be improved using specialized software, canonical and back-splice junction information can be extracted directly from BAM files or input separately as BED-formatted files containing the coordinates and quantities of each junction. The user is provided the flexibility of normalizing expression of each exon across all samples or for exon normalization to be confined within each sample (this helps visualize alternative splicing, intron retention, and exon exclusion). As introns are generally much larger than exons, an option to reduce intron size by a user-defined amount is also provided. In an effort to guide interpretation of gene specific splicing patterns, predicted or empirically determined RNA binding protein binding sites can be added to the plots (Figure 1B-C) by supplying a list of coordinates or utilizing the consensus binding sequences determined by *Ray et al*^*33*^. Because inverted ALU repeat elements impact RNA secondary structure, we have also incorporated the option to add a track of ALU elements to the plot.

**Figure 1.**
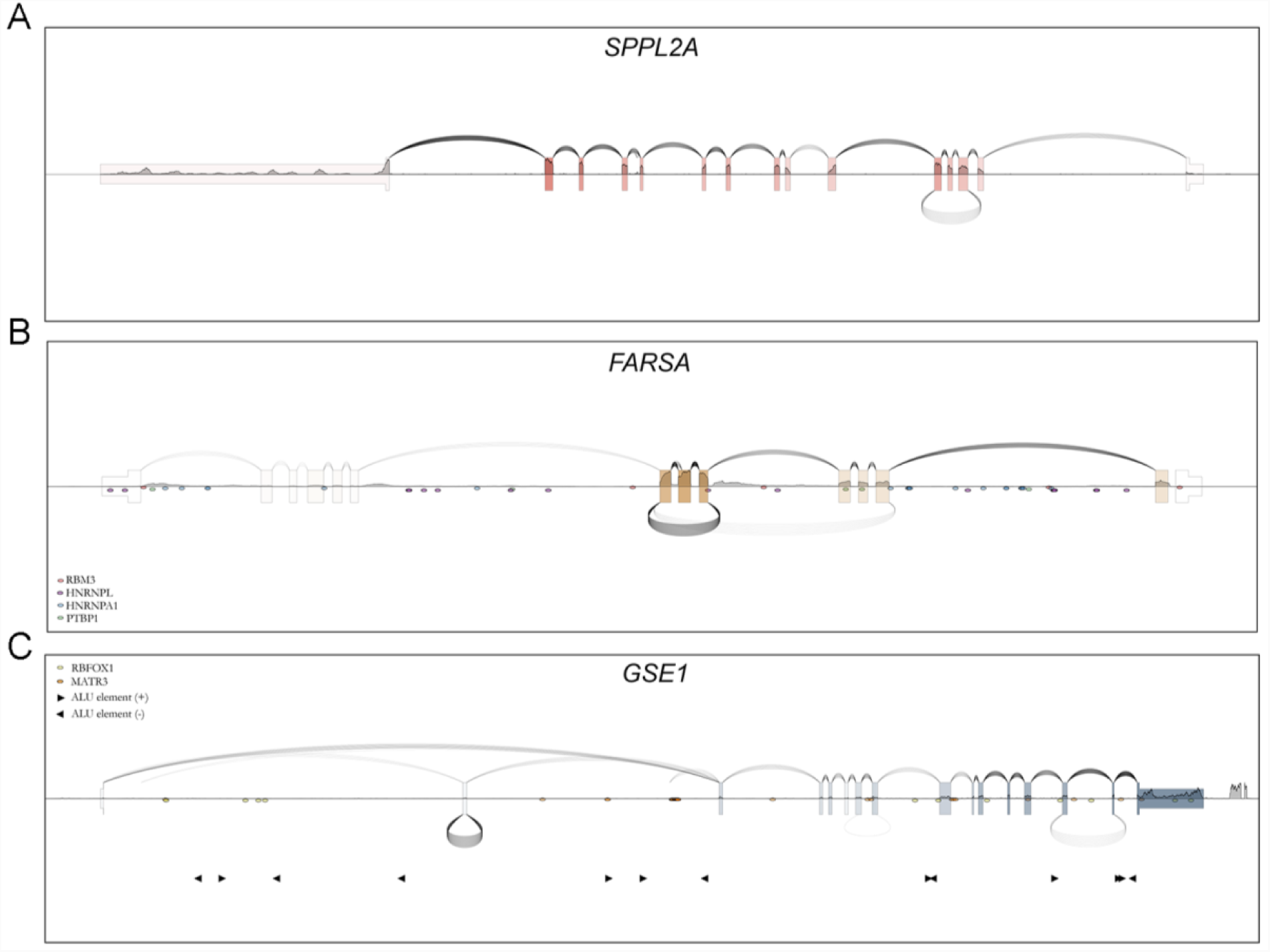
(A) SpliceV plot of SPPL2A expression in Akata cells. Coverage (exon level coverage; color intensity of each exon, single nucleotide level coverage; height of the horizontal line bisecting each exon) and forward splice junctions (arches above exons) was derived from sequencing a Poly-A selected library preparation. Back-splice junctions (curves below exons) were obtained from sequencing a ribodepleted, RNAse-R treated library preparation. (B) SpliceV plot of FARSA in Akata cells. All junctions and coverage were derived from a ribodepleted, RNAse-R treated sample. (C) SpliceV plot of GSE1 expression in ribodepleted, RNAse-R treated (back-splice junctions) and Poly-A selected (coverage and canonical junctions) SNU719 cells. Predicted binding sites for RNA binding proteins, RBM3, HNRNPL, HNRNPA1, PTBP1 are plotted along the FARSA transcript (B) and RBFOX1, and MATR3 sites are plotted along the GSE1 transcript (C). ALU elements are marked in (C).

## Results

Multiple computational pipelines have been developed to detect and quantify circRNAs from high throughput RNA sequencing data^18,21,34–36^. As circRNAs lack a poly-A tail, ribodepleted library preparations are essential for circRNA detection. RNA preparations can then be treated with the exonuclease, RNAse R, which exclusively digests linear RNAs, to increase the depth of circRNA coverage. To demonstrate the utility of SpliceV, we used libraries prepared from Poly-A selected (enriched for polyadenylated linear RNAs) or ribodepleted-RNase R-treated RNA from the Burkitt’s Lymphoma cell line, Akata, and the gastric carcinoma cell line, SNU719. Reads from each library were aligned using the STAR aligner v2.6.0a^37^ to generate BAM and splice junction BED files. We further processed our alignments using find_circ^21^ to interrogate the unmapped reads for back-splice junctions. Our first plot displays a prominent circular RNA formed via backsplicing from exon 5 to 3 of SPPL2A (Figure 1A). For this plot, backsplicing (under arches) derived from RNase R-seq data is plotted with forward splicing (over arches) and exon level (exon color intensity) and single nucleotide level (horizontal line graph) coverage from poly A-RNA-seq data from Akata cells to illustrate circRNA data in the context of linear polyA transcript expression. Exon level coverage display provides easy visualization of selective exon utilization: for example, using forward-and back-splicing and coverage data derived from RNase R-seq data (Akata cells) show enriched coverage of the circular RNA exons 6-8 of the FARSA gene (Figure 1B). Nevertheless, the simultaneous display of single nucleotide level coverage includes additional information that can help provide more detailed clarity in interpretation. For example, while the last exon of SPPL2A (Figure 1A) shows low exon level coverage, there is an evident drop in single nucleotide level coverage soon after the splice acceptor site, likely illustrating the utilization of an upstream polyA site (3’ UTR shortening^38^). Therefore, while exon level coverage provides illustrative qualities for some more macroscopic analyses (e.g. enriched exon coverage of circularized exons (Figure 1B in RNase R-seq data)), single nucleotide coverage provides granularity when needed.

The need that initially inspired us to develop SpliceV was the lack of available software to plot backsplicing in the context of coverage and forward splicing (for example, see Figure 1A). This is not only useful for simple presentation of circRNA splicing information, but can also aid interpretation. For example, the display of forward splicing and coverage from polyA-seq data in the context of backsplicing data from RNase R-seq data for the GSE1 gene provides evidence of circle formation of exon 2 which precludes its inclusion in the cognate linear GSE1 isoform (Figure 1C). In this case, exon 2 exclusion introduces a frameshift, ablating the canonical function of this gene.

To add utility to SpliceV in transcript biogenesis and isoform function analyses, we also incorporated the display of user defined RNA binding protein prediction (Figure 1B) and Alu repeats (Figure 1C). These features can assist the user in assessing the mechanisms of forward splicing, back splicing, alternative splicing, intron retention, etc. Further, since loaded RNA binding proteins control transcript localization as well as activity, these features can help assist the user in investigating transcript function.

To further illustrate the utility of SpliceV in investigational efforts, we next used SpliceV to visualize isoform level expression in two Gastric Carcinomas and one normal gastric tissue sample from The Cancer Genome Atlas (TCGA)^39^. Whole Exome Sequencing variant calls revealed that each of the two tumor samples had unique splice site mutations in the critical tumor suppressor gene, TP53^40^. The Genomic Data Commons pipeline^41^ for gene expression quantification revealed a slight increase in TP53 RNA levels in the tumor samples. Because the mutations in both tumors occurred in intronic regions, the impact on protein output is not easily determined. Using SpliceV to visualize RNA-seq data (Figure 2), however, revealed likely haplotypic ablation of the mutated splice acceptor (Figure 2B) or donor (Figure 2C) site in these two samples. This led to the utilization of cryptic splice sites that produced frameshifts in each of the resulting transcripts. Also evident in sample BR-8483, based on the single nucleotide coverage line graph, is extensive intron retention, likely causing the resulting intron retained transcript to be subjected to non-sense mediated RNA decay. In both of these cases, SpliceV was able to assist in determining the negative impact of these two mutations on TP53 function, findings that are otherwise opaque to the user.

**Figure 2.**
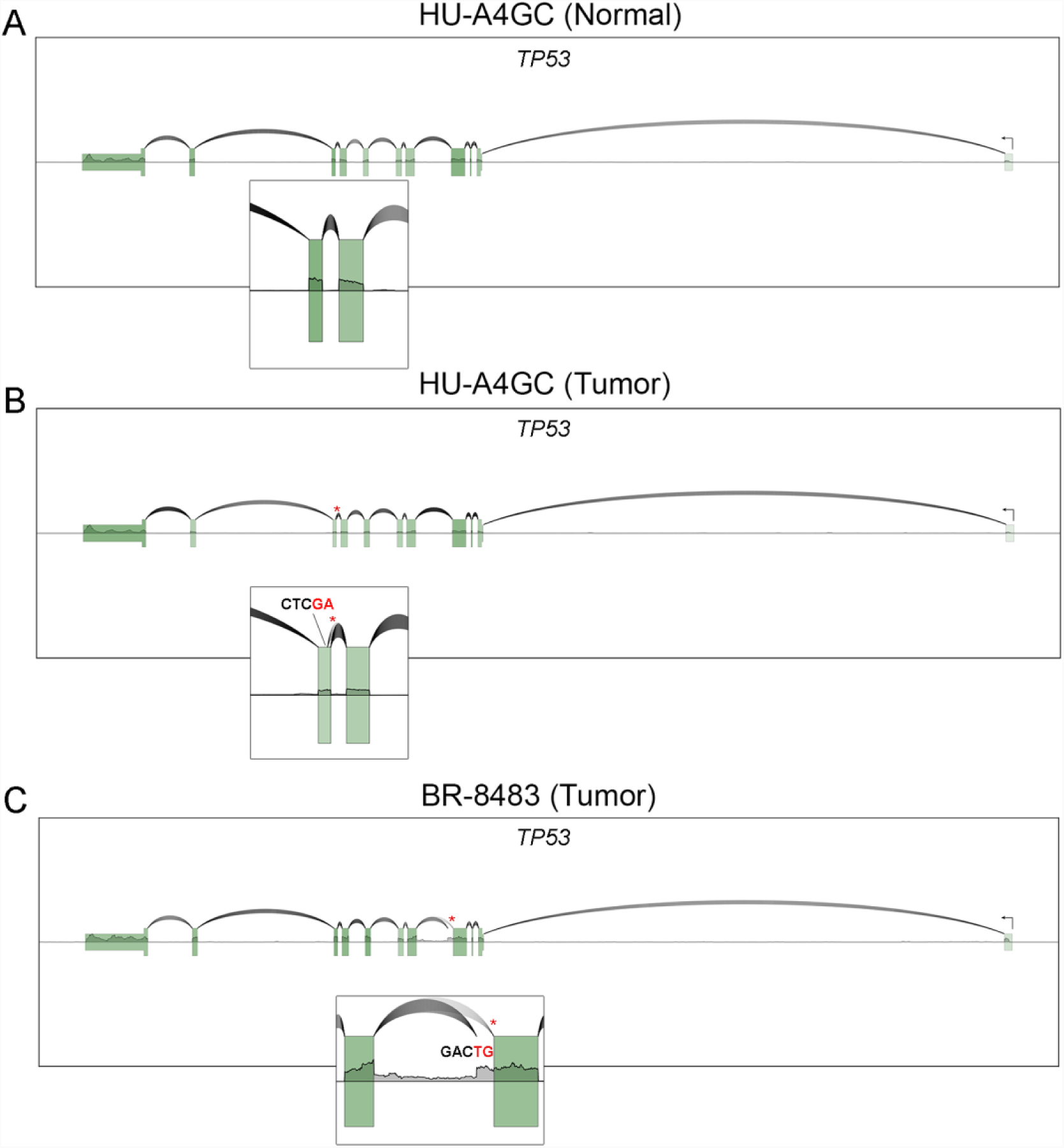
(A) SpliceV plot of RNA coverage and splicing from normal stomach tissue (wild type TP53). (B) SpliceV plot of a gastric tumor with a 1 base (T) deletion at a splice acceptor (chr17:7673610, HG38 genome build), disrupting the splicing from exon 8-9 and causing the utilization of a novel cryptic downstream splice acceptor at position chr17:7673590; resulting in a frameshift deletion. (C) SpliceV plot of a gastric tumor with a G>A splice donor variant at chr17:7675993. Part of the intron is retained (chr17:7675884-7675993) and a novel intronic splice donor site is utilized, with the same upstream acceptor. This introduces a frameshifting insertion into the protein coding sequence. Asterisks indicate the SNV location and insets are enlarged representations of the transcript structure. Nucleotide sequences at the cryptic splice sites are labeled, with the junctions occurring between the red and black bases in each figure. These samples were initially provided by The Cancer Genome Atlas^39^, with alignments obtained from the Genomic Data Commons^41^.

## Conclusion

Here we present a new tool, SpliceV, that facilitates investigations into transcript biogenesis, isoform function and the generation of publication quality figures for the RNA biologist. SpliceV is fast (taking full advantage of the random access nature of BAM files), customizable (allowing users to control plotting aesthetics), and can filter data and make cross-sample comparisons. It is modular in structure, allowing for the inclusion of new features in future package releases. SpliceV should provide value to the toolkit of investigators studying RNA biology and function and should speed the time frame from data acquisition, data analysis to publication of results.

## Acknowledgements

This work was supported by the National Institutes of health grants, R01AI106676, and P01CA214091 and the Department of Defense grant, W81XWH-16-1-0318. The funders had no role in study design, data collection and analysis, decision to publish, or preparation of the manuscript

