## Supplemental File 1 for "SpliceV: Analysis and publication quality printing of linear and circular RNA splicing, expression and regulation"

Usage:

*splicev -b [one or more bam files] -gtf [one gtf file] -g [gene to plot]*

all arguments:

**-h, --help** show help message

**-b, --bam** Path to bam file(s)

**-t, --transcript** Name of transcript to plot (must match "transcript_id" field of gtf file)

**-g, --gene** Name of gene to plot (overrides "-t" flag). Will plot the longest transcript derived from that gene

**-gtf** Path to gtf file

**-bsj** Path to backsplice junction bed-formatted files (listed in the same order as the input bam files)

**-sj** Path to canonical splice junction bed-formatted files (listed in the same order as the input bam files)

**-stranded** If strand-specific sequencing, indicate 'forward' if upstream reads are forward strand, otherwise indicate 'reverse' (True-Seq is 'reverse').

**-is, --intron-scale** The factor by which intron white space should be reduced

**-c, --color** Exon color. Hex colors (i.e. "\#4286f4". For hex, an escape "\" must precede the argument), RGB (i.e. 211,19,23) or names (i.e. "red")

**-f, --filter** Filter out splice junctions and circles that have fewer than this number of counts.

**-n, --normalize** Normalize coverage between samples

**-rc, --reduce_canonical** Factor by which to reduce canonical curves

**-rbs, --reduce_backsplice** Factor by which to reduce backsplice curves

**-ro, --repress_open** Do not automatically open plot in browser

**-en, --exon_numbering** Label exons

**-rnabp** List of RNA binding proteins to plot.

**-fa** Path to fasta file

**-format** Output image format {SVG,PDF,PNG,JPG,TIFF}

**-alu** Path to Alu bed file
